# SKP2-mediated FBXO2 proteasomal degradation drives hepatocellular carcinoma progression via stabilizing Hsp47

**DOI:** 10.1101/2024.03.28.586926

**Authors:** Cailin Xue, Fei Yang, Guojian Bao, Jiawu Yan, Rao Fu, Minglu Zhang, Jialu Ding, Jiale Feng, Jianbo Han, Xihu Qin, Hua Su, Beicheng Sun

## Abstract

Accumulating studies highlight that dysregulated E3 ubiquitin ligases are associated with the onset and advancement of cancers. Nevertheless, the impact and mechanism of most E3 ubiquitin ligases on tumorigenesis and tumor metastasis remain poorly understood. Here, we show that loss of FBXO2, an E3 ubiquitin ligase, accelerates hepatocellular carcinoma (HCC) tumor growth and metastasis to the lung through stabilizing heat shock protein 47 (Hsp47). Downregulation of FBXO2, caused by DNA-PKcs-medicated phosphorylation at serine 17 and E3 ligase SKP2-mediated ubiquitination at lysine 79 and subsequent proteasomal degradation, is observed in tumor tissues compared to their parallel non-tumor tissues resected from patients with HCC. Patients whose tumors are enriched for SKP2 or Hsp47 or express low levels of FBXO2 have poor median survival compared to those whose tumors have reversed levels of SKP2, FBXO2 and Hsp47. Together, FBXO2 acts as a tumor suppressor in HCC development. The components of the SKP2-FBXO2-Hsp47 axis provide newly prognostic and therapeutic factors for anti-HCC.

## Introduction

Hepatocellular carcinoma (HCC) is one of the most common and highly aggressive digestive system cancers worldwide with a 5-year survival rate of less than 20% at advanced stage in China^1,2^. According to the latest statistics in 2019, HCC ranks fourth in incidence, with approximately 450,000 new cases, and fifth in mortality, with approximately 5,000 deaths. Prevalence of metastases is the leading cause of poor outcome^3^. However, the mechanisms by which HCC tumors metastasize to other organs remain largely unknown.

Ubiquitin-proteasome system (UPS), the major proteolytic system that controls protein degradation, has been found to perform important role in cancer development^4^. The UPS is composed of two parts, ubiquitin ligases and 26S proteasomes^5^. While the former is responsible for ubiquitin conjugation to the substrate through a multistep cascade reaction involved the E1, E2, and E3 ubiquitin ligase enzymes, the latter is capable of cleaving ubiquitin-tagging substrates^6^. E3 ubiquitin ligases, which facilitate attachment of ubiquitin onto particular proteins via their lysine residues, have been found to display double roles in cancer progression serving as either tumor promoters or suppressors^7,8^. In light of approval from the U.S. Food and Drug Administration (FDA) regarding use of Bortezomib-a proteasome inhibitor-against multiple myeloma and mantle cell lymphoma^9^, researchers have focused greater interest towards developing compounds that target E3 ligases due to their higher selectivity when compared to proteasome inhibitors whose efficacy ranges broadly yet often come coupled with side effects.

E3 ligases are classified into three types, homologous to the E6AP carboxyl terminus (HECT) type, Really interesting new gene (RING)-finger type, and the U-box type. The SKP1-cullin1-F-box protein (SCF) E3 ligase complex, also known as cullin-RING ligase 1 (CRL1), which belongs to the RING-finger type, is the largest family of E3 ubiquitin ligases^10^. It consists of an adaptor protein SKP1, a scaffold protein Cullin-1 (CUL1), a RING protein RBX1 or RBX2, and a substrate-recognizing F-box protein whose type determines the specificity of substrate recognition. Out of F-box proteins, FBXO2, also known as Fbx2, Fbs1, OCP1 and NFB42, is a poorly characterized and special one that ubiquitinates uniquely to N-glycosylated proteins^11^. FBXO2 is known to promote the development of endometrial cancer, gastric cancer and osteosarcoma through distinct cancer hallmark pathways^12-14^, suggesting a diversity of substrates of FBXO2 in various cancers. However, the impact and mechanism of FBXO2 on HCC progression remain unknown.

Extracellular matrix (ECM) produced by cancer cells serves as metastatic niches to facilitate cancer cell colonization in the distant organs^15^. Heat shock protein 47 (Hsp47), a collagen-specific chaperone assisting collagen secretion and deposition, has been found to be highly expressed during epithelial mesenchymal transformation (EMT) and promote metastasis^14^. However, the regulation of Hsp47 expression and its role in promoting metastasis are still unknown.

Here we found that FBXO2 is downregulated in human HCC tumors compared to its parallel non-tumor tissues. Inhibition of FBXO2 via SKP2-mediated ubiquitous degradation or through its loss-of-function variant (FBXO2 W78C) promotes HCC tumor growth and metastasis through stabilizing Hsp47, suggesting FBXO2 is a tumor suppressor of HCC. Patients with SKP2^high^, FBXO2^low^, or Hsp47^high^ tumors have a considerably worse median survival than do patients with reversed expression of these markers.

## Results

### FBXO2 is an HCC tumor suppressor

To discover the aberrantly expressed E3 ubiquitin ligases in HCC tumors, a high-throughput RNA sequencing of four pairs of human HCC tumor tissues and their adjacent non-tumor tissues was performed. Bioinformatic analysis revealed that the mRNA level of the E3 ligase FBXO2 is upregulated in HCC tumor tissues compared to adjacent non-tumor tissues (Extended Data Fig. 1a), which was confirmed by real-time polymerase chain reaction (RT-PCR) analysis of FBXO2 mRNA in 24 pairs of human HCC tumor tissues and adjacent non-tumor tissues. FBXO2 mRNA was upregulated in most (16/24) tumor tissues (Extended Data Fig. 1b). Unexpectedly, immunohistochemical (IHC) analysis of 258 pairs of human HCC tumors and para-tumor tissues revealed that FBXO2 protein level is frequently downregulated in tumor tissues (149/258, *P*<0.001) compared to para-tumor tissues (Fig. 1a, b). Similar results were obtained in liver tissues resected from a mouse model of HCC induced by N-nitrosodiethylamine (DEN) and carbon tetrachloride (CCl_4_) (Fig. 1c). Discrepancy between mRNA and protein abundance of FBXO2 implies that the diminished expression of FBXW2 protein in HCC tumors is post-transcriptional.

**Fig. 1.**
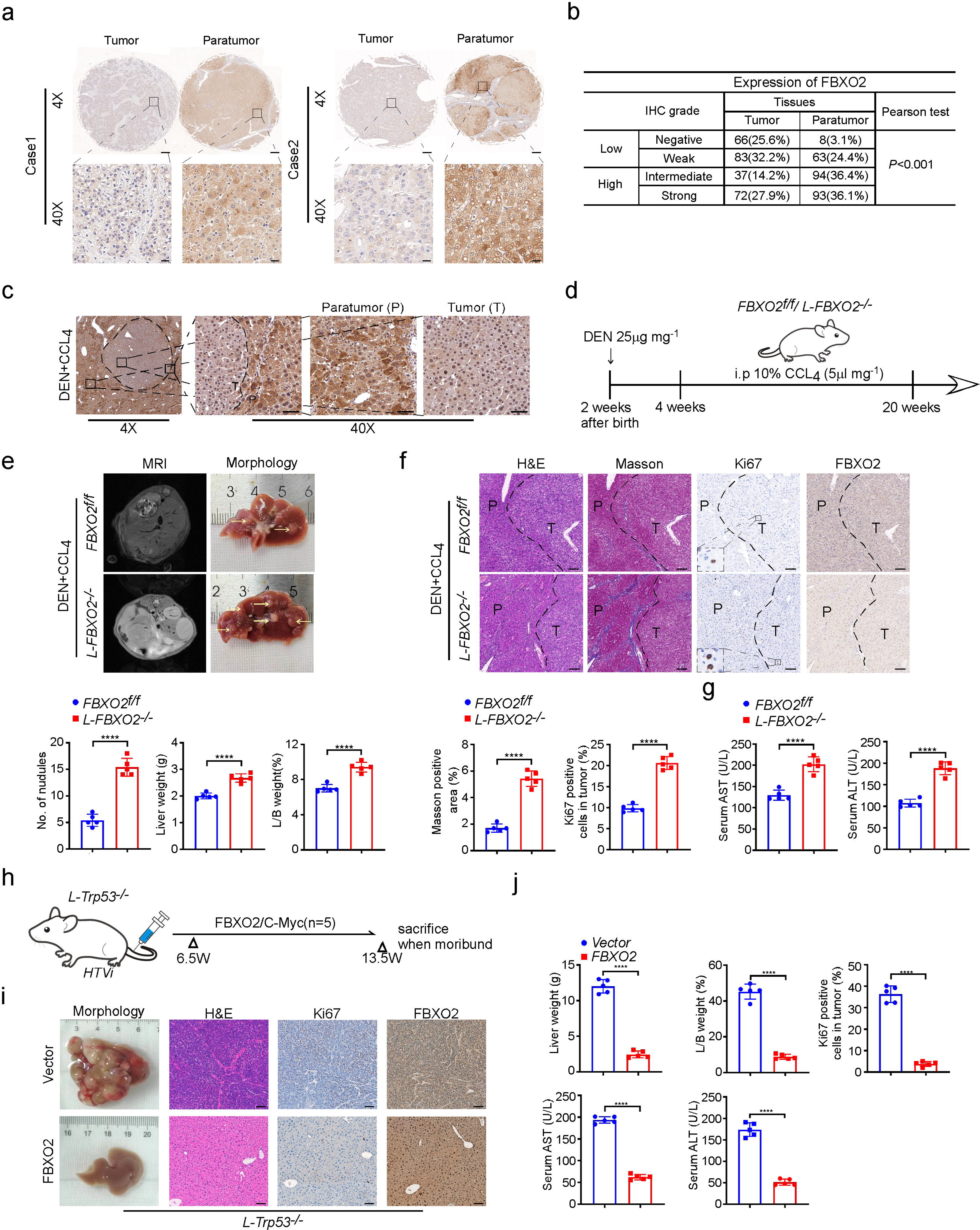
Ablation of FBXO2 accelerates hepatocarcinogenesis. **a**, Representative immunohistochemistry (IHC) of 258 pairs of resected human hepatocellular carcinoma (HCC) and adjacent non-tumorous tissues. Upper scale bar, 200 μm, lower scale bar, 20 μm **b**, Numbers and percentages of above human HCC specimens positive for FBXO2 protein (arbitrarily indicated as negative, weak, intermediate or strong). The correlation between FBXO2 and Hsp47 was analyzed by a two-tailed chi-square test. *n* = 258, \*\*\**P* < 0.001. **c,** Representative IHC images of FBXO2 in liver tissues derived from the DEN/CCL_4_-induced HCC mice. Scale bar, 50 μm. **d,** Diagram of the DEN/CCl_4_-induced HCC protocol in wild-type (*FBXO2^f/f^*) mice and mice whose hepatocytes are deleted with FBXO2 via Albumin (Alb)-cre (*L-FBXO2^-/-^*). **e-g,** Heavier liver tumor burdens in DEN/CCl_4_-treated *L-FBXO2^-/-^* mice compared to DEN/CCl_4_-treated *FBXO2^f/f^* mice indicated by representative images of liver MRI, morphology, tumor number, liver weight and liver/body ratio (**e**), and H&E, Masson, Ki67 and FBXO2 staining of liver sections as well as quantification of positive Masson and Ki67 staining (**f**) and serum AST and ALT levels **(g)**. Scale bar, 100 μm (*n*=5). Data are displayed as the mean ± SD, \*\*\*\**P*<0.0001. **h,** Diagram of hydrodynamic tail vein injection of plasmid expressing C-Myc to induce HCC in hepatocyte-Trp53 knock out mice(*L-Trp53^-/-^*). **i, j,** FBXO2 overexpression inhibits C-Myc-induced HCC in *L-Trp53^-/-^* mice indicated by H&E, Ki67 and FBXO2 staining (**i**) of liver sections as well as the quantification of liver weight, liver/body weight (L/B) ratio, Ki67 positive cells, and serum AST and ALT levels (**j**). Scale bar, 100 μm (*n*=5). Data are displayed as the mean ± SD. Statistical significance was determined by a two-tailed *t* test. \*\*\*\**P*<0.0001

To investigate the effect of FBXO2 downregulation on HCC tumor progression, we first generated hepatocyte-specific deletion of *FBXO2 (L-FBXO2^-/-^)* mice by crossing *FBXO2^Flox/Flox^ (FBXO2^f/f^)* mice with *Alb-Cre* mice. Ablation of FBXO2 in hepatocytes was verified by PCR and immunoblot (IB) analysis (Extended Data Fig. 1c, d). Then *FBXO2^f/f^* mice and *L-FBXO2^-/-^* mice were intraperitoneally administrated with one dose of 25 μg mg^-1^ DEN and 10% CCl_4_ (5μl mg^-1^) for 20 weeks to induce tumorigenesis of HCC (Fig. 1d). The liver resected from *L-FBXO2^-/-^* mice showed a much heavier tumor burden than the liver from *FBXO2^f/f^* mice after treatment of DEN and CCl4 indicated by magnetic resonance imaging (MRI), liver morphology, increased tumor number and liver weight as wells as liver to body (L/B) weight ratio of *L-FBXO2^-/-^* mice (Fig. 1e). This was confirmed by increased fibrosis indicated by masson staining, and higher expression of proliferation marker (Ki67) in FBXO2-ablated tumors (Fig. 1f). As expected, significantly increased serum aspartate aminotransferase (AST) and alanine transaminase (ALT) were observed in *L-FBXO2^-/-^*mice, indicating exacerbated liver damage (Fig. 1g). Similar results were observed in another mouse model of HCC, induced through overexpression of c-Met and β-catenin in the liver of *FBXO2^f/f^* and *L-FBXO2^-/-^* mice via hydrodynamic injection and a sleeping beauty transposase system (Extended Data Fig.1e-g). FBXO2 deletion accelerated c-Met and β-catenin-induced HCC tumor growth. Reversely, overexpression of FBXO2 dramatically repressed c-Myc-driven tumorigenesis in mouse liver with *Trp53-*deleted hepatocytes (*L-Trp53^-/-^*), or inhibited the growth of tumor cells returned to the liver through splenic injection of (cell lines) in nude mice (Fig. 1h-j, Extended Data Fig.1h, i). These results suggest that FBXO2 acts as a tumor suppressor of HCC.

### FBXO2 suppresses HCC metastasis in vitro and in vivo

Metastasis remains the major obstacle towards effective therapy against HCC. To investigate whether FBXO2 effects in HCC metastasis, we first measured FBXO2 protein expression in one normal human hepatocyte line L02 and multiple human HCC cell lines (Huh7, HCC-LM3, MHCC-97H, Hep3B and HepG2). Compared to L02 cells, HepG2, Huh7, and Hep3B cells exhibited a higher, whereas MHCC-97H showed an equal and HCC-LM3 showed a lower protein level of FBXO2 (Fig. 2a). Then we overexpressed or ablated FBXO2 in Hep3B cells with a moderate high FBXO2 expression, or in HCC-LM3 cells with a low FBXO2 expression, and investigated their abilities of migration, invasion and metastasis. While ablation of FBXO2 dramatically enhanced migration and invasion of Hep3B cells indicated by the assays of transwell migration, invasion and wound-healing (Fig. 2b-d), overexpression of FBXO2 performed an opposite effect on them in Hep3B and HCC-LM3 cells (Fig. 2e, f and Extended Data Fig. 2a-c), suggesting an inhibitory effect of FBXO2 on metastasis. In order to verify it in vivo, parental or FBXO2-ablated Hep3B cells stably expressing luciferase were transplanted into nude mice through tail vein injection to investigate tumorigenesis in lung. As expected, loss of FBXO2 significantly enhanced Hep3B cell growth in lung indicated by luciferase activity, lung morphology, tumor number, and H&E staining (Fig. 2g-i). These results suggest that FBXO2 inhibits HCC metastasis to lung.

**Fig. 2.**
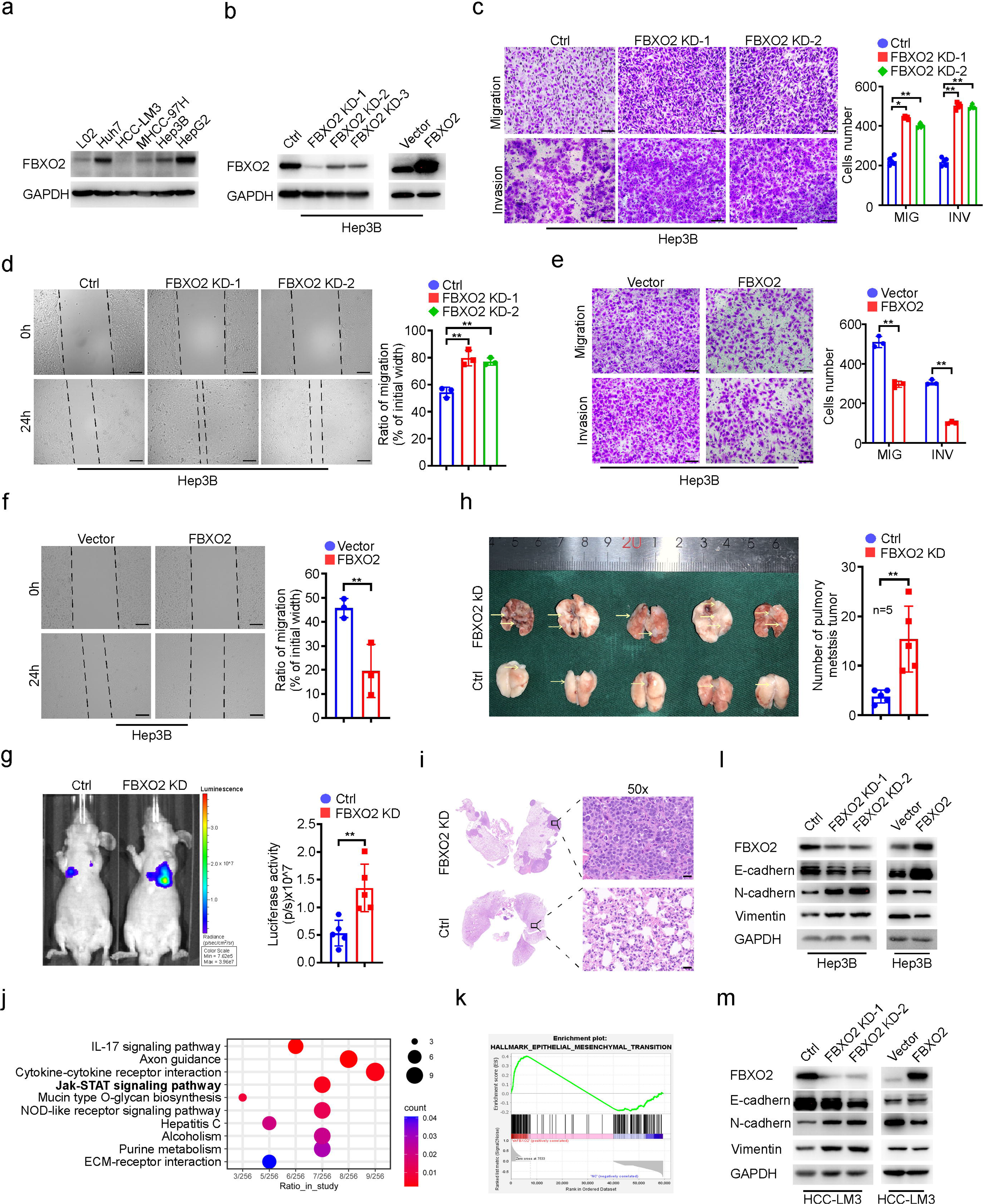
FBXO2 suppresses HCC tumor cell migration, invasion, and metastasis to lung. **a,** Immunoblot (IB) analysis of the indicated proteins in a normal human hepatocyte cell line (L02) and human HCC cell lines (Huh7, HCC-LM3, HCC-97H, Hep3B, and HepG2). **b,** IB analysis the indicated proteins in FBXO2-knockdown (KD) or -overexpression Hep3B cells. **c, d,** FBXO2 KD promotes migration or invasion of Hep3B cells via transwell chamber migration and invasion assays (**c**) and wound-healing assay (**d**). Scale bar, 50 μm. Data are displayed as the mean ± SD. Statistical significance was determined by a two-tailed *t* test. \**P* < 0.05, \*\**P* < 0.01. **e, f,** FBXO2 overexpression inhibits migration or invasion of Hep3B cells through transwell chamber migration and invasion assays (**e**) and wound healing assay (**f**). Scale bar, 50 μm. Data are displayed as the mean ± SD. Statistical significance was determined by a two-tailed *t* test. \*\**P* < 0.01. **g,** Representative images and quantification of the bioluminescence signal of pulmonary metastases eight weeks after tail vein injection of parental (Ctrl) or FBXO2 KD Hep3B cells into nude mice. Data are displayed as the mean ± SD (*n* = 5). Statistical significance was determined by a two-tailed *t* test. \*\**P* < 0.01. **h,** Gross images of lung morphology and quantification of metastatic tumors in lung from above mice. Data are displayed as the mean ± SD (*n* = 5). Statistical significance was determined by a two-tailed *t* test. \*\**P* < 0.01. **i,** H&E staining of lung sections from the above mice. Boxed areas are further magnified. Scale bar, 20 μm. **j,** Top 10 enriched KEGG pathways of RNA-seq data from parental (shNC) and FBXO2-KD (shFBXO2) Hep3B cells. **k,** Gene set enrichment analysis (GSEA) of epithelial-mesenchymal transition gene sets in the expression profiles of above cells. **l, m,** IB analysis of the indicated proteins in Hep3B cells (**l**) or HCC-LM3 cells (**m**) with or without FBXO2 KD or overexpression.

To understand the underlying mechanism for enhanced metastatic ability of FBXO2-ablated HCC cells, we performed RNA-seq in parental and FBXO2-ablated Hep3B cells. Gene set enrichment analysis (GSEA) and Kyoto Encyclopedia of Genes and Genomes (KEGG) pathway enrichment analysis showed that FBXO2 deletion upregulates JAK-STAT signaling pathway which is a cancer intrinsic driver of tumor growth and metastasis (Fig. 2j, k). Consistently, FBXO2 ablation triggers the expression of mesenchymal cell markers vimentin and N-cadherin, and represses the expression of epithelial marker E-cadherin, whereas FBXO2 overexpression showed an opposite effect on the expression of these makers (Fig. 2l, m), suggesting an inhibitory role of FBXO2 in EMT. These results suggest that loss of FBXO2 promotes HCC metastasis through upregulation of EMT.

### FBXO2 interacts with Hsp47 and promotes its ubiquitination and subsequent proteasomal degradation

To investigate how FBXO2 inhibits EMT, we performed affinity purification-mass spectrometry (MS) of the interaction proteins with FBXO2, resulting Hsp47 (SERPINH1) which was found to promote EMT shows a strong affinity with FBXO2 indicated by the relatively high score (Fig. 3a, Supplementary Table 4). Bioinformatic analysis of the MS data indicated that Hsp47 is a functionally related protein of FBXO2 (Fig. 3b). Immunoprecipitation assay revealed that Hsp47 is able to bind to FBXO2 in HEK293, Hep3B and HCC-LM3 cells (Fig. 3c, d), whereas another functional-related protein EGFR is unable (Fig. 3d), indicating a specific interaction between FBXO2 and Hsp47. FBXO2 has been found to interact specially with glycosylated proteins through Tyr-278 (Y278) and Trp279 (W279) at its FBA domain which locates in the C-terminal of F-box proteins and is responsible for specific substrate recognition^16,17^. To identify whether Y278 and W279 are required for FBXO2 and Hsp47 interaction, we made multiple variants of FBXO2 and detected their affinities with Hsp47. As expected, while deletion of F-box domain (Flag-FBXO2-C), which behaved as parental FBXO2, did not affect FBXO2 and Hs47 binding, absence of FBA domain (Flag-FBXO2-N) or the site mutations whose Y278 or W279 was changed to alanine (Y278A or W279A) buried their abilities to bind to Hsp47 (Extended Data Fig. 3a).

**Fig. 3.**
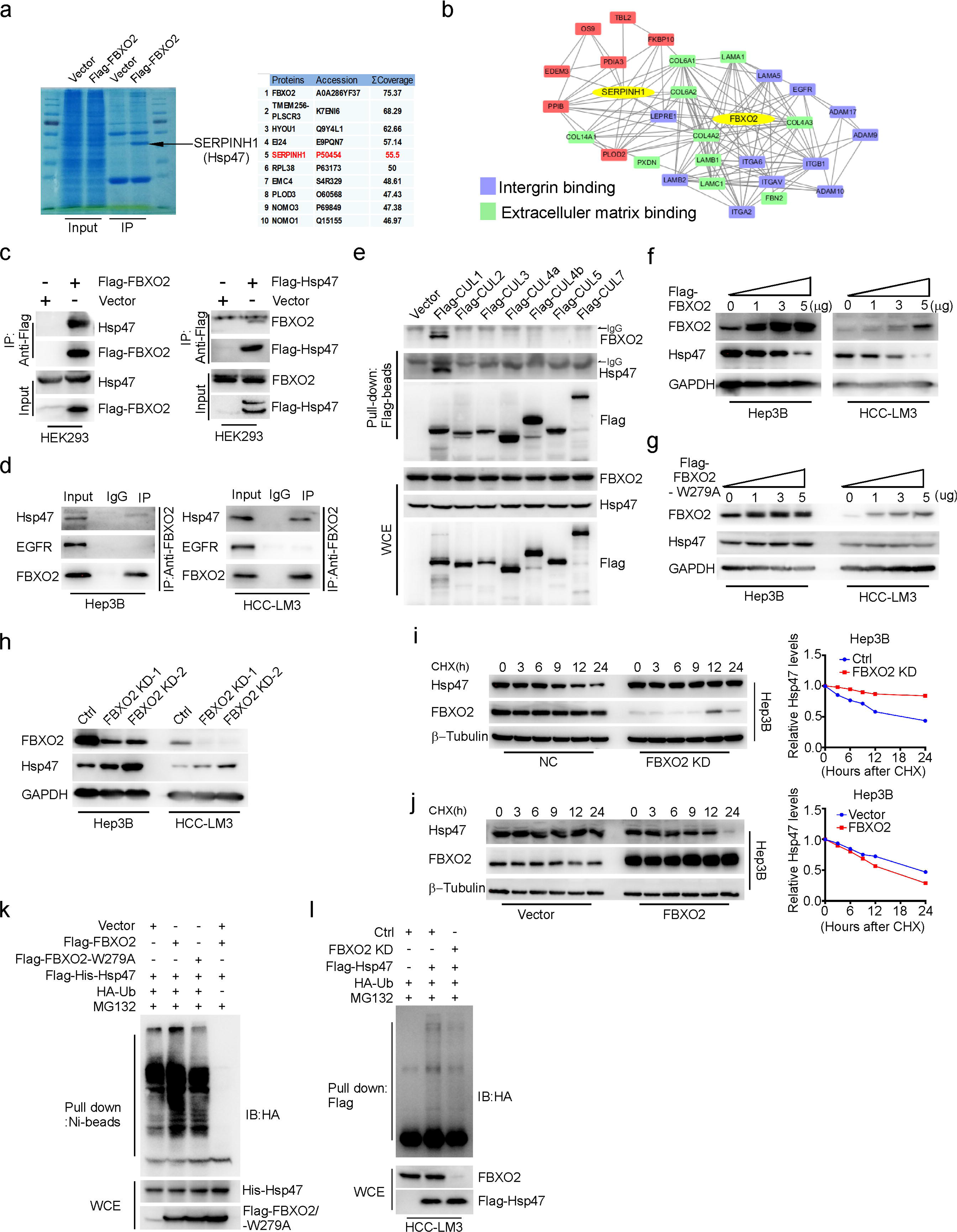
FBXO2 ubiquitylates Hsp47 and shortens its half-life. **a,** LCLMS/MS analysis of FBXO2 binding proteins. **b,** Bioinformatics analysis of the functional correlation between FBXO2 and Hsp47 based on MS data. **c**, Co-immunoprecipitation (Co-IP) analysis of the interaction between FBXO2 and Hsp47 in MG132 treated-HEK293 cells transfected with or without Flag-tagged FBXO2 or Hsp47. IP was carried out with anti-Flag followed by MG132 (10 μm) treatment for 6 hours and the precipitates were examined by IB with the indicated antibodies. **d**, Co-IP of endogenous Hsp47 with FBXO2 in Hep3B and HCC-LM3 cells. FBXO2 was immunoprecipitated, and the precipitates were analyzed using anti-Hsp47 and anti-EGFR. **e**, Co-IP of FBXO2 and Hsp47 with the Flag-tagged indicated proteins in transfected HEK293 cells. IP was carried out with anti-Flag, and the precipitates were immunoblotted with the indicated antibodies. **f, g** IB analysis of the indicated proteins in Hep3B and HCC-LM3 cells overexpressing different amounts of Flag-tagged FBXO2 (**f**) or FBXO2-W279A (**g**). **h,** IB analysis of the indicated proteins in parental (Ctrl) and FBXO2 KD Hep3B or HCC-LM3 cells. **i, j,** IB analysis of the indicated proteins in Hep3B cells with or without KD (**i**) or overexpression (**j**) of FBXO2 followed by cycloheximide (CHX, 50μg ml^-1^) treatment for the indicated time periods. While FBXO2 ablation extends the half-life of Hsp47 (**i**), FBXO2 overexpression shortens it (**j**). The band intensity was quantified using Image J, normalized to β-Tubulin first, then normalized to the t=0 time point. **k,** FBXO2 promotes Hsp47 ubiquitylation: IB analysis of Ni-bead pull-down and whole cell extraction (WCE) derived from HEK293 cells transfected with the indicated constructs. **l**, Ablation of FBXO2 inhibits Hsp47 ubiquitylation: IB analysis of Flag tag pull-down and WCE derived from parental or FBXO2-ablated HCC-LM3 cells transfected with the indicated constructs followed by treatment of MG132.

The F-box proteins commonly assemble into SCF complex which consists of Cullin 1 (CUL1), SKP1, RBX1 and a variable F-box protein to perform its catalytic activity^18^. We found that FBXO2 and Hsp47 specially bind to CUL1 but not any other CUL proteins by a pull-down assay using Flag-beads (Fig. 3e), which is consistent with the previous results showed that FBXO2 is a component of SCF complex^16,19^. In addition, overexpression of parental FBXO2 but not FBXO2-W279A decreased Hsp47 proteins in a dose-dependent manner, whereas ablation of FBXO2 effected as the proteasome inhibitor MG132 increased them (Fig. 3f-h and Extended Data Fig. 3b). Given that NEDD8-activating enzyme (NAE) catalyzed-CUL neddylation is required for SCF E3 ligase activity, we blocked CUL neddylation by a NAE inhibitor MLN4924 and found MLN4924 triggered Hsp47 accumulation (Extended Data Fig. 3c). Notably, both MG132 and MLN4924 are also able to increase FBXO2 protein level (Extended Data Fig. 3b, c), which is consistent with our former results showed FBXO2 expression is posttranscriptional regulated in human HCC specimens (Fig. 1a, and Extended Data Fig. 1a, b), suggesting FBXO2 expression may be controlled by ubiquitination-mediated proteasomal degradation. Hsp47 mRNA level was not affected by genetic manipulations of FBXO2 (Extended Data Fig. 3d). Further we found inhibition of SCF E3 ligase by ablation of FBXO2 or CUL1, or administration of MLN4924 remarkably extended the protein half-lives of Hsp47, whereas FBXO2 overexpression reversed them (Fig. 3i, j, Extended Data Fig. 3e-h). Moreover, parental FBXO2 other than FBXO2-W279A promoted the K48-linked ubiquitination of Hsp47 (Fig. 3k, Extended Data Fig. 3i), whereas FBXO2 ablation inhibited it (Fig. 3l). Altogether, these results suggest that FBXO2 promotes Hsp47 ubiquitination and subsequent proteasomal degradation.

### FBXO2 depletion promotes HCC metastasis to lung by stabilizing Hsp47

To investigate the biological significance of FBXO2-mediated Hsp47 degradation in HCC progression, we examined its effect on migration, invasion and metastasis of HCC cell lines through manipulation the expression levels of FBXO2 and Hsp47. Loss of FBXO2 increased, or overexpression of FBXO2 decreased tumor cell migration and invasion, whereas Hsp47 ablation or overexpression reversed them separately (Fig. 4a, b and Extended Data Fig. 4a, b). Accordingly, Hsp47 deletion reversed FBXO2 ablation-induced downregulation of E-cadherin and upregulation of N-cadherin and Vimentin., Similarly, Hsp47 overexpressing cells showed opposite expression levels of EMT-related proteins to FBXO2- overexpressing cells (Fig. 4c and Extended Data Fig. 4c, d). Consistently, Hsp47 ablation blocked FBXO2 deletion-induced HCC cell growth in lung indicated by luciferase activity, tumor numbers and HE staining of lung tissue sections (Fig. 4d-f). These results suggest that loss of FBXO2 promotes HCC metastasis to lung though stabilizing Hsp47.

**Fig. 4.**
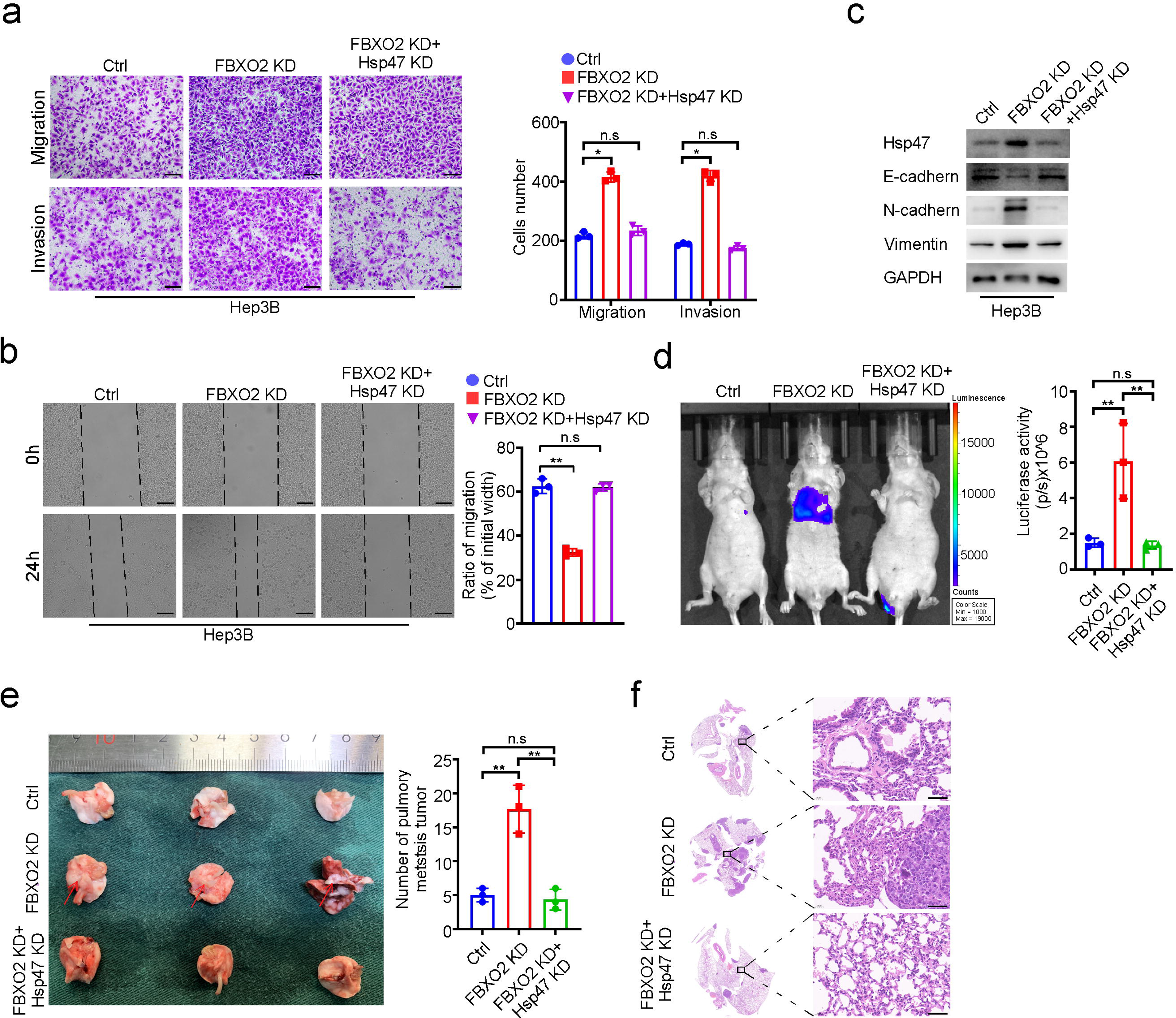
FBXO2 depletion promotes HCC metastasis to lung by stabilizing Hsp47. **a, b,** Representative images and quantification of the effects of FBXO2 KD or double (FBXO2 and Hsp47) KD on the migration and invasion of Hep3B cells through transwell chamber migration and invasion assays **(a)** and wound healing assay (**b**). Scale bar, 50 μm. **c**, IB analysis of the indicated proteins in the above cells. **d,** Representative images and quantification of the bioluminescence signal of pulmonary metastases eight weeks after tail vein injection of parental (Ctrl), FBXO2 KD, or FBXO2 and Hsp47 double KD Hep3B cells into nude mice. **e,** Gross images and quantification of tumor numbers of lung resected from above mice. **f,** H&E staining of lung sections from the above mice. Boxed areas were further magnified. Scale bar, 50 μm. Results in (a, b, d, e) (nL=L3), are mean±SD. Statistical significance determined by two-tailed t-test. *PL<L0.05, **PL<L0.01.

### The E3 ligase SKP2-mediated FBXO2 ubiquitination relies on DNA-dependent protein kinase activity

As mentioned above, FBXO2 expression is probable controlled by ubiquitination-mediated proteasomal degradation in human HCC specimens. To verify this, we first investigated which F-box protein can bind to FBXO2 through a pull-down assay using Flag beads in HEK293 cells expressing different flag-tagged F-box proteins and found only Flag-tagged SKP2 is able to bind to FBXO2 (Fig. 5a). Similar results were observed by IP assays showing either exogenous or endogenous FBXO2 can bind to endogenous SKP2 (Fig. 5b, c). SKP2 ablation increased FBXO2 protein levels and extends FBXO2 half-life, whereas ectopically expressed SKP2 decreased them (Fig. 5d-g). Meanwhile, FBXO2 mRNA level was not affected by SKP2 expression levels (Extended Data Fig. 5a, b). Further we found SKP2 promotes FBXO2 polyubiquitylation via K48 linkage (Extended Data Fig. 5c, d). Importantly, MS analysis of the ubiquitylated FBXO2 suggested the potential ubiquitylation site of FBXO2 by SKP2 locates at K79 (Fig. 5h). To confirm this, Flag-tagged FBXO2 variant in which the lysine residue at 79 was changed to alanine via site-directed mutagenesis was constructed. Ubiquitylation assessment affirmed that SKP2 fails to ubiquitinate FBXO2-K79A (Fig. 5i). These results suggest that SKP2 acts as a ubiquitin ligase that promotes ubiquitylation of FBXO2 at K79 and its subsequent degradation.

**Fig. 5.**
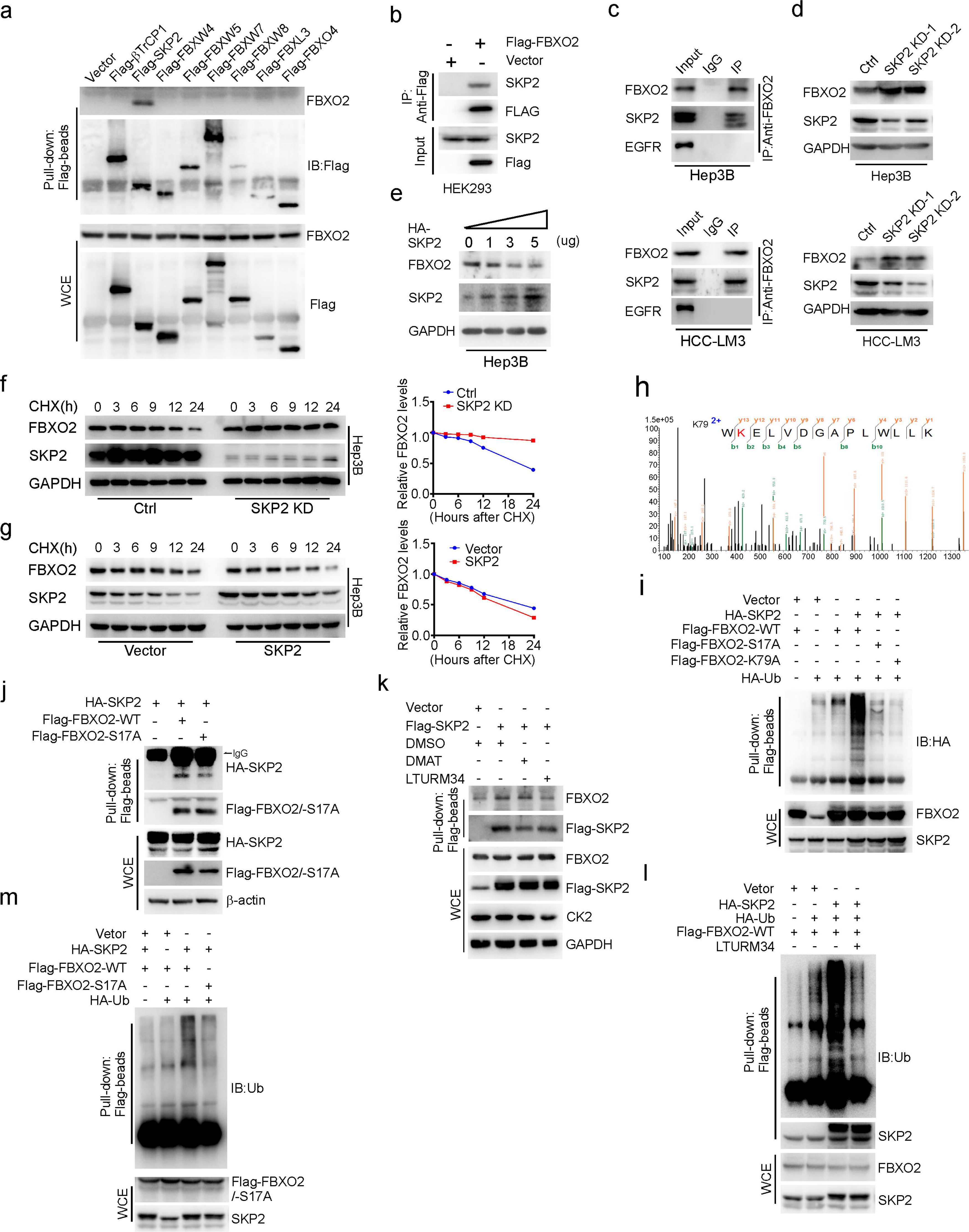
SKP2 shortens the FBXO2 half-life by ubiquitinating it for degradation. **a,** Pull-down assay of FBXO2 with the indicated Flag-tagged proteins in transfected HEK293 cells. Pull-down was carried out with Flag-beads, and the precipitates were immunoblotted with the indicated antibodies. **b**, Co-IP of SKP2 with FBXO2 in HEK293 cells transfected with the indicated constructs. **c**, Co-IP of SKP2 with FBXO2 in Hep3B and HCC-LM3 cells. IP was carried out with anti-FBXO2. **d,** IB analysis of the indicated proteins in Hep3B and HCC-LM3 cells with or without SKP2 ablation. **e,** IB analysis of the indicated proteins in Hep3B cells transfected with the indicated amount of Flag-tagged SKP2 vector. **f**, **g,** IB analysis of the indicated proteins in Hep3B cells with or without ablation (**f**) or overexpression (**g**) of SKP2 followed by CHX (50μg ml^-1^) treatment for the indicated periods. The band density was quantified using Image J, normalized to GAPDH first, then normalized to the t=0 time point. **h,** The annotated MS/MS spectrum of the tryptic peptide showing ubiquitylation at residue K79 of FBXO2. **i,** SKP2 promotes polyubiquitination of FBXO2-WT but not its mutant FBXO2-S17A or FBXO2-K79A. IB analysis of Flag-beads pull-down and WCE derived from HEK293 cells transfected with the indicated constructs. **j**, Pull-down of HA-tagged SKP2 with Flag-tagged FBXO2 or FBXO2-S17A in HEK293 cells. Precipitates and WCE were immunoblotted with the indicated antibodies. **k,** IB assay of Flag-beads pull-down and WCE derived from HEK293 cells with or without Flag-tagged SKP2 transfection followed by treatment of DMSO, CK2 inhibitor (DMAT, 10 μM) or DNA-PKcs inhibitor (LTURM34, 10 μM) for 24 h. **l**, LTURM34 inhibits SKP2-mediated ubiquitination of FBXO2: IB analysis of Flag-beads pull-down and WCE derived from the indicated construct-transfected HEK293 cells treated with or without LTURM34.

It is known that in most cases, phosphorylation is prerequisite for a substrate to bind to a F-box protein for targeted ubiquitylation and degradation by the SCF ubiquitin ligases^20^. To examine whether the interaction between SKP2 and FBXO2 is phosphorylation dependent, we constructed a FBXO2 phosphorylation-dead mutant (S17A) on SKP2 degron motif which is highly conserved in FBXO2 protein sequence among different species (Extended Data Fig. 5e). SKP2 failed to shorten the half-life of FBXO2-S17A and ubiquitinate it (Fig. 5i and Extended Data Fig. 5f). Indeed, FBXO2-S17A performed a lower affinity with SKP2 compared to parental FBXO2 (FBXO2-WT) (Fig. 5j). These results imply SKP2-mediated FBXO2 ubiquitination requires phosphorylation of FBXO2 at S17. To verify this, we forecasted the potential kinase of FBXO2 at S17 through SCANSITE 4.0 (https://scansite4.mit.edu/4.0/#home) and found that DNA-PKcs (DNA-dependent protein kinase catalytic subunit) and CK2 (casein kinase 2) were the two most likely kinases of FBXO2 indicated by the high score (Extended Data Fig. 5g). CK2 inhibitor DMAT did not affect FBXO2 and SKP2 interaction, whereas inhibition of DNA-PKcs via its inhibitor LTURM34 or siRNA dramatically inhibited their interaction (Fig. 5k and Extended Data Fig. 5h). Consistently, suppression of DNA-PKcs prevented SKP2-mediated ubiquitination and half-life shortening of FBXO2 (Fig. 5l and Extended Data Fig. 5j). Taken together, these results suggest that FBXO2 phosphorylated by DNA-PKcs is prerequisite for its subsequent ubiquitylation via SKP2.

### Low levels of SKP2 or Hsp47, or high levels of FBXO2 correlate with improved survival

IHC of surgically resected human HCC showed that around 58% (149/258) tumors contained low amounts of FBXO2 and most of them exhibited higher levels of staining for SKP2 (89/149) and Hsp47 (96/149) than did FBXO2^high^ tumors (Fig. 6a, b), suggesting a negative correlation between SKP2 and FBXO2, or between FBXO2 and Hsp47. In addition, low FBXO2 expression correlated with a more advanced tumor AJCC stage (*P*=0.043) and Edmonds stages (*P=*0.036) in patients (Fig. 6c), implying a poor prognosis. Notably, patients with SKP2^high^ or FBXO2^low^, or Hsp47^high^ tumors had a considerably worse median survival than did patients with reversed expression of these markers (Fig. 6d-f). These results are consistent with those obtained in our preclinical HCC models, suggesting that SKP2-mediated FBXO2 degradation promotes Hsp47 accumulation which drives HCC progression.

**Fig 6.**
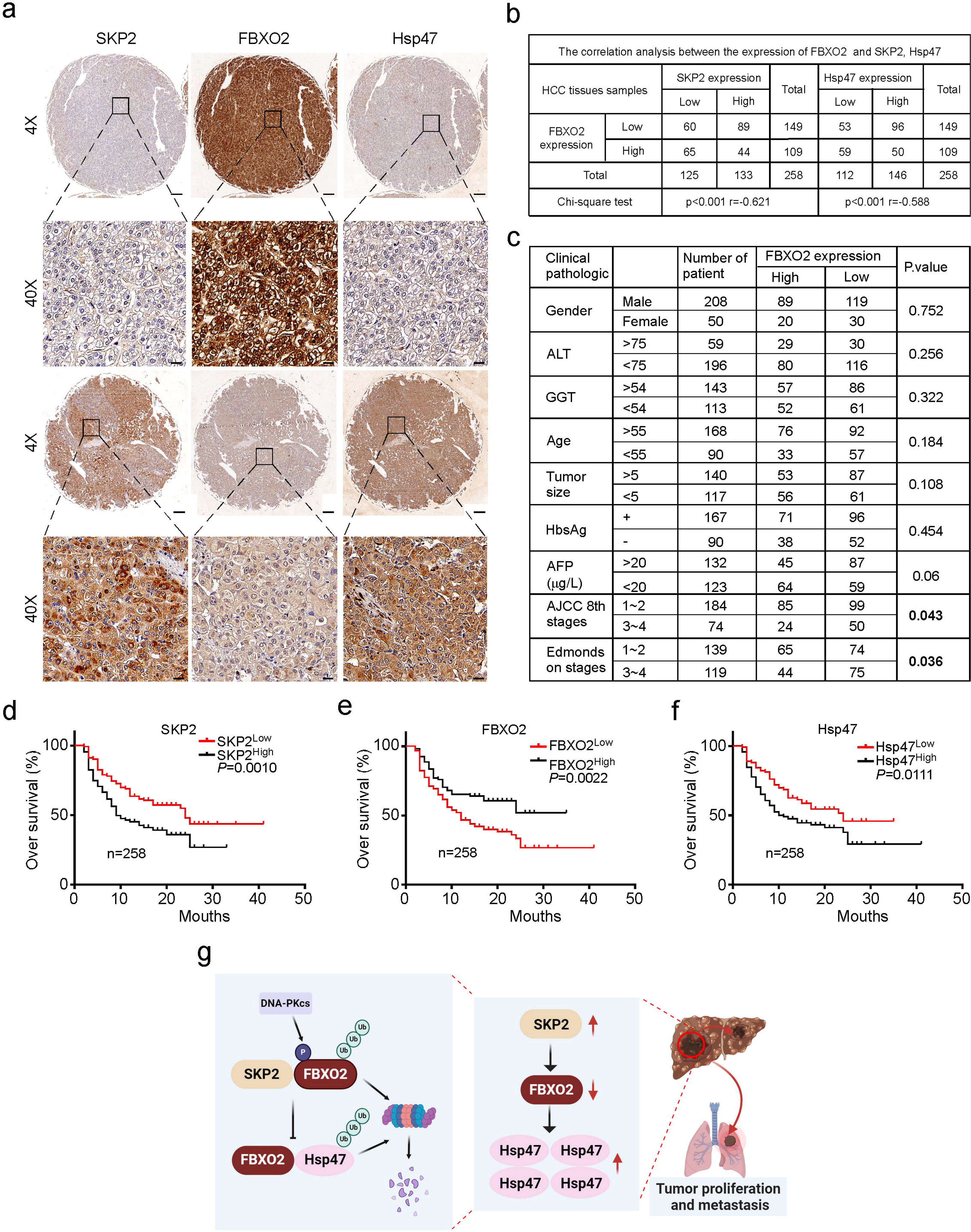
High SKP2, low FBXO2 or high Hsp47 expression predicts poor survival of patients with HCC. **a,** IHC analysis of the indicated proteins in human HCC specimens. Scale bar (4X), 200 μm, scale bar (40X), 20 μm. **b,** Correlation of expression levels between FBXO2 and SKP2 in above tissues analyzed by a two-tailed Chi-square test. **c,** Clinicopathological characteristics of the investigated HCC patients in **a** and **b** (*n*=258). **d-f,** Comparisons of overall survival between patients stratified according to the expression levels of SKP2 (**d**), FBXO2 (**e**) or Hsp47. **g,** Schematic depiction of the mechanism underlying FBXO2-mediated HCC progression.

## Discussion

Although E3 ubiquitin ligase plays crucial roles in tumorigenesis and metastasis, our understanding still remains limited compared towards plethora associated functions across various types of E3 ubiquitin ligases involving neoplastic transformation process. The E3 ligase FBXO2 has been previously found to act as a tumor promoter in multiple cancers^12-14^ For example, FBXO2 targeted SUN2 or Fibrillin1 (FBN1) for ubiquitination and degradation to promote ovarian cancer or endometrial carcinoma development separately^12^. FBXO2 also promoted osteosarcoma progression by activating the STAT3 signaling pathway^14^. Here we found FBXO2 acts as a tumor suppressor of HCC via degrading Hsp47. These results suggest that in distinct tumor contexts, FBXO2 displays opposite effects on tumor progression via various substrates preference. The successful treatments thus require understanding the effects of specific E3 ligase on each cancer other than one-size-fits-all effects. Emerging techniques encompassing structurally defined mechanisms responsible for substrate recognition mediated via monovalent molecular glues and bivalent proteolysis-targeting chimeras^21^, or the multiplex CRISPR screening platform to assign E3 ligases to their cognate substrates at scale will facilitate to complete this outstanding question in the near future^22^. Our results showed that loss of FBXO2 promotes both tumorigenesis and metastasis. Although Hsp47 has been known to promote EMT, how it enhances cell proliferation is unclear. One possibility is increased tumor cell-derived collagen I homotrimers which effect as soluble collagens (cleaved collagens) promote tumor metabolism as Hsp47 is a molecular chaperone of collagens to facilitate their synthesis and secretion^23-27^. Although various substrates of FBXO2 have been identified in different cancers^12-14^, it was previously unknown that how FBXO2 expression is regulated during cancer development. Here we found FBXO2 protein level is controlled by the E3 ligase SKP2, which depends on DNA-PKcs-mediated FBXO2 phosphorylation. These results are consistent with the previous studies showed that SKP2^28^ or DNA-PKcs accelerates HCC progression^29^. SKP2 was found to promote cell proliferation, migration and invasion though AMPK and CRAM1-induced autophagy^28-30^. These results indicate that the E3 ligase is able to integrate multiple pathways via ubiquitinating different substrates to regulate cancer development. DNA-PKcs was known to facilitate tumor growth via upregulation of DNA repair^31,32^. Our results emphasize the perspective that DNA-PKcs is not limited to fixing DNA breaks but also effects in other cellular functions^33^. A deeper understanding of how the E3 ligase integrates different oncogenic pathways and how DNA-PKcs activity is regulated in various cellular processes is another area deserves priority research.

## Acknowledgments

This work was supported by grants from the National Natural Science Foundation of China (81930086 and 82120108012 to B.S.; 82002931 and 82372644 to F.Y.; 82372884 to H.S.), the Project of Innovation & Entrepreneurship, Jiangsu Province (No. 2020-30136), the Clinical Research Special Project of Anhui Provincial Department of Science and Technology (202204295107020008 to B.S.), the Research Program of Anhui Provincial Department of Education (2022AH010070 to B.S.), and the Shanghai Municipal Committee of Science and Technology (23ZR1413600 to H.S.).

## Author Contributions

C.X. performed most experiments; C.X. and B.G. performed qPCR, animal work and analyzed data; C.X., J.Y, R.F, M.Z. and J.D. performed IB analysis; J.F and H.S. drew the summary chart; B.S., H.S., F.Y., and X.Q. conceived designed, and supervised the study; H.S., F.Y., B.S. and C.X. wrote the manuscript, with all authors contributing and providing feedback and advice.

## Methods

### Tissue samples

A total of 258 cases of HCC and corresponding adjacent non-tumor tissues were obtained from Nanjing Drum Tower Hospital from September 2018 to September 2020. None of the patients received radiotherapy or chemotherapy before surgery. Written informed consent was obtained from all patients involved in this study. All experimental procedures were approved by the Ethics Committee of Nanjing Drum Tower Hospital and met the guidelines of the Declaration of Helsinki and International Ethical Guidelines for Biomedical Research Involving Human Subjects. The clinicopathologic characteristics of HCC patients involved in the research are listed in Figure 6c.

### Cell culture

All cell lines, including L02, Hep3B, HCC-LM3, HepG2, Huh7, and MHCC-97H, were cultured in Dulbecco’s modified Eagle’s medium (DMEM) (Invitrogen, USA) containing 10% fetal bovine serum (FBS) (Gibco, USA). and 1% penicillin streptomycin (Invitrogen, USA) and maintained in a humidified incubator at 37°C with 5% CO_2_. All cells were purchased from the Type Culture Collection of the Chinese Academy of Sciences (Shanghai, China). All the cells were routinely tested and were negative for mycoplasma.

### Plasmids, siRNA and lentiviral infection

The human full-length cDNAs of *FBXO2* and *Hsp47* with Flag tags were synthesized and subcloned into the lentiviral vector pCDH-CMV-MCS-EF1-Puro by Sangon Biotech (Sangon Biotech, China). The plasmids, including Flag-βTrCP1, Flag-SKP2, Flag-FBXW2, Flag-FBXW4, Flag-FBXW5, Flag-FBXW7, Flag-FBXW8, Flag-CUL1, Flag-CUL2, Flag-CUL3, Flag-CUL4a, Flag-CUL4b, Flag-CUL5, Flag-CUL7, and Flag-CUL8, were preserved in our laboratory. The variants of FBXO2 or Hsp47 were generated by PCR using the ClonExpress II One Step Clony Kit (Vazyme, China) according to the instructions. For pT3-EF1α−FBXO2 plasmids matched with the sleeping beauty system, EcoRI (NEB, USA) and BsmI (NEB, USA) were used to cut the sequence, and the cDNA of FBXO2 was cloned into pT3-EF1α−MCS with the ClonExpress II One Step Cloning Kit (Vazyme, China).

The siRNAs targeting the sequences of human *FBXO2*, *Hsp47*, *SKP2* and *DNA-PKcs* were purchased from GenePharma (Shanghai, China). The DNA fragments of the shRNA targets human *FBOX2*, *Hsp47*, and *SKP2* were synthesized by GenScript (GenScript, China) and subcloned into the lentiviral vector pLKO.1-puro (Addgene, USA). All plasmids used in this study were verified by DNA sequencing. The sequences of the siRNA and shRNA are listed in Supplementary Table 1. For siRNA or plasmid transient transfection, Lipo3000 (Invitrogen, USA) was used according to the manufacturer’s protocols. For lentivirus packaging, the targeted plasmid and virus packaging plasmids (psPAX2, pMD2.G) were co-transfected into 293T cells. To establish stable cell lines, the cells were infected with the lentivirus expressing shRNA or cDNA in the presence of polybrene (5 μg/mL) (Sigma, USA), followed by selection with puromycin (5 μg/mL) (Sigma, USA) for 2 weeks.

### Migration and invasion assays

In migration and invasion assays, Corning Costar Transwell 24-well plates (8 μm) (Corning, USA) were used to evaluate the migration and invasion ability of human HCC cell lines. For migration assays, a total of 10^5^ cells were resuspended in 200 μl serum-free medium and plated into the upper chambers, and 800 μl DMEM containing 10% FBS was added to the lower chambers. For migration assays, a total of 5ⅹ10^4^ cells were seeded into upper chambers. For invasion assays, a total of 10^5^ cells were seeded into upper chambers precoated with 30 µg Matrigel (BD Biosciences, USA). The cells were incubated at 37°C with 5% CO_2_ for 12-18 h. The non-migrated cells in the upper chamber were removed, and the cells that migrated onto the bottom were fixed with methanol and stained with1% crystal violet at the indicated time in the figures. The measurements were done on randomly-selected fields of view.

### Wound healing assays

Cells were plated on a six-well plate. A 10-μL plastic pipette tip was used to create a scratch wound when the cell confluence exceeded 90%. Then, the cells were washed with PBS twice and maintained in DMEM with 1% FBS. The scratch wounds were photographed at 0 and 24 h after wounding. The scratch healing area was analyzed by Image J software.

### In vivo lung metastasis assay

For lung metastasis assays, a total of 3 × 10^6^ human HCC tumor cells were resuspended in 200 μL PBS and injected into 4-week-old BALB/c nude mice through the tail vein. After 8 weeks, the mice were injected with luciferin at 7.5Lg/kg to monitor metastases using luminescence imaging system (Perkin-Elmer, USA). Then, the mice were sacrificed, and the lungs of the nude mice were fixed in 4% paraformaldehyde, embedded in paraffin, and stained with hematoxylin-eosin (H&E). All animal study procedures were approved by the Animal Care Committee of Nanjing Drum Tower Hospital and followed the National Institutes of Health Animal Care and Use Guidelines.

### Chemicals, and inhibitors

Puromycin and proteasome inhibitor MG132 were purchased from MedChemExpress (MedChemExpress, USA). LTURM34 (DNA-PK inhibitor), KU-55933 (ATM kinase inhibitor), MK-2206 (AKT1 inhibitor) and CHX (cycloheximide) were purchased from Selleck (Selleck, USA).

### Quantitative real-time PCR (qRT-PCR)

Total RNA was extracted using TRIzol (Invitrogen, USA). HiScript Q RT SuperMix (Vazyme,China) were used for reverse transcription following the manufacturer’s protocol. qRT-PCR was conducted on ABI Prism 7900HT (Applied Biosystems, USA) using ChamQ Universal SYBR qPCR Master Mix (Vazyme, China) according to the manufacturer’s instructions. Relative expression levels of target genes were normalized against the level of β-actin or GAPDH. The primers used in this study were synthesized by GenScript (GenScript, China). The primers are listed in Supplementary Table 2.

### Immunoblot (IB) and antibody

For IB, the tissues or cells were harvested and lysed in RIPA buffer (Beyotime, China) supplemented with 1% protease inhibitors and phosphatase inhibitors. Protein concentration was measured using a bicinchoninic assay (BCA) kit (Beyotime, China). The protein was resolved on SDS-polyacrylamide gels, and then transferred to polyvinylidene difluoride (PVDF) membranes. The PVDF (Millipore, German) membranes were then incubated in 5% milk at room temperature for 1 h, followed by staining with primary antibody at 4°C overnight. Then, the PVDF membranes were incubated with anti-mouse or anti-rabbit HRP- conjugated antibodies for 2 h, and developed with chemiluminescence (ECL) (Tanon, China). Immunoreactive bands were detected by Tanon Imager. The antibodies used in this study are listed in Supplementary Table 3.

### Co-Immunoprecipitation (co-IP)

The cells were treated with MG132 (MCE, USA) (10 μM) for 6 h before lysed in IP lysis buffer (50 mM Tris-HCl, pH 8.0, 120 mM NaCl, 0.5% NP40, 1 mM EDTA) containing 1% protease inhibitor and phosphatase inhibitors. Then IP was performed using the beads-conjugated anti-Flag antibody. Generally, 2 μg of antibody were added to 1 ml of cell lysate and incubated at 4°C overnight. The beads were then washed with IP lysis four times, and supplemented with 25 µl 2 × protein loading buffer. Subsequently, the beads were heated and the supernatant were resolved by SDS-PAGE, and analyzed by IB.

### Mass spectrometric analysis

For mass spectrometric analysis, the HEK-293 cells were transfected with Flag-FBXO2 plasmids for 3 days. The cells were then harvested and lysed followed by IP. Flag-FBXO2 protein complexes were visualized with Coomassie brilliant blue staining. The Coomassie brilliant blue staining bands were analyzed by LC-MS/MS using a FAMOS autosampler. The National Center for Biotechnology Information nonredundant protein sequence database was used to analysis the sample spectra. The identified proteins were listed in Supplementary Table 4.

### In vivo ubiquitination assay

To investigate the ubiquitination of Hsp47 by FBXO2, Flag-Hsp47, HA-Ub, and FBXO2 or FBXO2-S17A were co-transfected into HEK293 cells. To investigate the ubiquitination of FBXO2 by SKP2, Flag-FBXO2, HA-Ub, and SKP2 were co-transfected into HEK293 cells. To identify the ubiquitin linkage in the Hsp47 polyubiquitination chain, HA-Ub (wild-type), HA-Ub-K48, HA-Ub-K63, HA-Ub-K48R, and HA-Ub-K63R were transfected into HEK293 cells along with Flag-Hsp47 and FBXO2. To determine the type of ubiquitin linkage in the FBXO2 poly-ubiquitylation chain, HA-Ub (wild-type), HA-Ub-K48, HA-Ub-K63, HA-Ub-K48R, and HA-Ub-K63R were transfected into HEK293 cells along with Flag-FBXO2 and SKP2. Cells were lysed followed by IP analysis using beads-conjugated anti-Flag antibody. The Hsp47-poly-HA-Ub and FBXO2-poly-HA-Ub were detected by IB using anti-HA antibody.

### Half-life assay

Cells were treated with CHX (Selleck, USA) 50 μg ml^-1^ and harvested at the indicated time. Then, the cells were lysed followed by IB. The densitometry of relative Hsp47 and FBXO2 levels was quantified using Image J software.

### Immunohistochemical assay

Paraffin-embedded tissue sections were dewaxed with xylene and rehydrated through graded ethanol. Then, antigen retrieval was conducted at 95°C with 0.1% sodium citrate buffer (pH 6.0) for 20 min, followed by incubation with 3% H_2_O_2_ and 1% bovine serum albumin-PBS for blocking the non-specific protein binding. Then, the tissue sections were incubated with the indicated primary antibody overnight at 4°C. After washing with PBS for three times, the tissues were incubated with HRP-conjugated secondary antibody for 30 min at room temperature and stained with a DAB substrate kit (Vetcorlabs, USA). The slides were photographed under light microscope. Immunohistochemical staining was semi-quantitated using a modified labeling score system (H score), based on the intensity of the staining and the percentage of the stained area. The staining intensity scores were divided into four levels (0, negative; 1, weak; 2, moderate; 3, strong). H score got a range of 0-300 by multiplying the staining intensity scores and the percentage of the positive area. The comprehensive scores of 0 ∼ 5, 5 ∼ 100, 101 ∼ 200, 201 ∼ 300 were divided into negative, weak, medium, and strong expression level. The diagnoses of all samples were confirmed by histological review. Each sample was evaluated blind by three pathologists, with the average score as the final score.

### Mice

C57BL/6 background *FBXO2^Flox/Flox^(FBXO2^f/f^)* and BALB/c nude mice were obtained from GemPharmatech (GemPharmatech, China). *Trp53^Flox/Flox^(Trp53^f/f^)* mice were a gift from the Laboratory Animal Center of Southern Medical University. To generate hepatocytes-FBXO2 knockout mice (*L-FBXO2^-/-^*), we crossed *FBXO2^f/f^* mice with Alb-Cre mice. *Trp53^Flox/Flox^ (Trp53^f/f^)* mice were crossed with Alb-Cre mice to obtain hepatocytes-*Trp53* knockout mice (*L-Trp53^-/-^*). Primer for PCR genotyping are listed in Supplementary Table 5. All mice were fed in a 12-h light-dark room and had free access to water and food.

### Mouse HCC models

For the chemical HCC model, *FBXO2^f/f^* and *L-FBXO2^-/-^* littermates were injected intraperitoneally with N-nitrosodiethylamine (DEN) (25 mg/kg) at 2 weeks postnatal, followed by weekly intraperitoneal injection of carbon tetrachloride (CCL_4_) (0.5 mL/kg, dissolved in olive oil). Mice were sacrificed at 5 months old, and the livers were collected for further analysis.

To establish an β-catenin/c-Met-induced HCC model, we hydrodynamically injected β-catenin/C-Met plasmid cDNA into 6.5-week-old male mice through the tail veil. Each mouse received a total of 22.5 mg cDNA, 10 mg pT3-EF1α-ΔN90-β-catenin (Human), 10 mg pT3-EF1α-c-Met (Human) and 2.5 mg pCMV-SB100, encoding the sleeping beauty transposase and transposon. To construct c-Myc-induced HCC in *L-Trp53^-/-^* mice, we hydrodynamically injected 15 mg pT3-EF1α-FBXO2 (Human)10 mg or pT3-EF1α, as a control; 5 mg pT3-EF1α−c-Myc (Human) and 2 mg pCMV-SB100 into each mouse though tail vein. The mice were monitored by palpation, and were sacrificed when the tumor burden was heavy. Eight weeks after the injection, the mice were euthanized. The plasmids, including pT3-EF1α-ΔN90-β-catenin and pT3-EF1α-c-Met, were a gift from the Xin Chen’s lab at University of California, San Francisco. All animal experiments were approved by the Animal Ethics Committee of The Affiliated Drum Tower Hospital of Nanjing University Medical School.

### Statistical analysis

Student’s *t* test was used to analyze the significance of the difference between two quantitative variables. One-way ANOVA was used to compare the differences among multiple groups. The Pearson correlation test was used to analyze the correlation between the expression of two proteins in HCC. Kaplan-Meier survival curves were analyzed by log rank test. All statistical analyses were performed by SPSS 22.0 software, and \**P* < 0.05, \*\**P* < 0.01, and \*\*\**P* < 0.001 were considered significant.

## Figure legends

**Extended Data Fig. 1.**
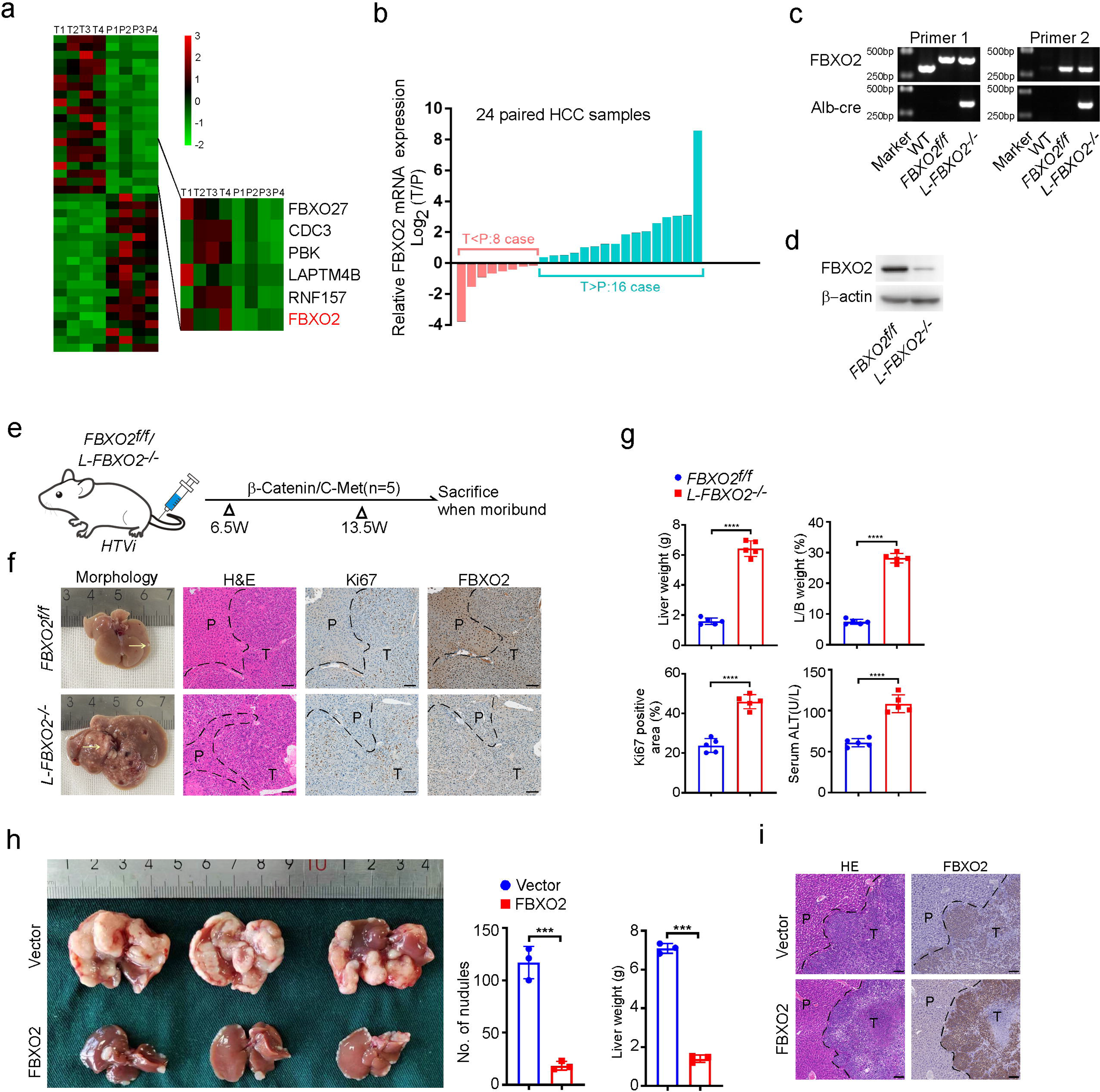
FBXO2 inhibits hepatocarcinogenesis. **a,** Clustered heatmap of the differentially expressed mRNAs in four paired human HCC tumor tissues and adjacent non-tumorous tissues. Upregulated genes are highlighted in red, and downregulated genes are in green. Fold changes >5; \**P* < 0.05. **b,** qRT-PCR analysis of Hsp47 mRNA in 24 paired human HCC tumor and adjacent non-tumorous tissues. **c**, PCR analysis of genomic DNA isolated from the tails of the indicated mice. **d**. Immunoblot (IB) analysis of FBXO2 expression in total liver cells derived from the indicated mice. **e**, Diagram of hydrodynamic tail vein injection of the plasmid expressing β-catenin/C-Met to induce-induced HCC in the indicated mice. 6.5-week-old *L-FBXO2^-/-^ /FBXO2^f/f^*male mice were hydrodynamically injected withβ-catenin/C-Met plasmid and sacrificed 7 weeks post injection. **f, g**, Heavier HCC tumor burdens were observed in β-catenin/c-Met-treated *L-FBXO2^-/-^* mice compared to *FBXO2^f/f^* mice indicated by representative images of liver morphology, and H&E, Ki67 and FBXO2 staining of liver sections (**f**) as well as quantification of liver weight, L/B ratio, Ki67 positive cells and serum ALT levels (**g**). Scale bar, 100 μm. Data are displayed as the mean ± SD. Statistical significance was determined by a two-tailed *t* test. \*\*\*\**P*<0.0001. **h,** Representative gross images of liver resected from the mice splenic injected with parental (vector) or FBXO2-overexpressing Hep3B cells. Quantification of tumor number and liver weight was shown on the right. Data are displayed as the mean ± SD (*n*=3). Statistical significance was determined by a two-tailed *t* test. \*\*\**P*<0.001. **(i)**H&E and FBXO2 staining of liver sections from above mice. Scale bar, 100 μm.

**Extended Data Fig. 2.**
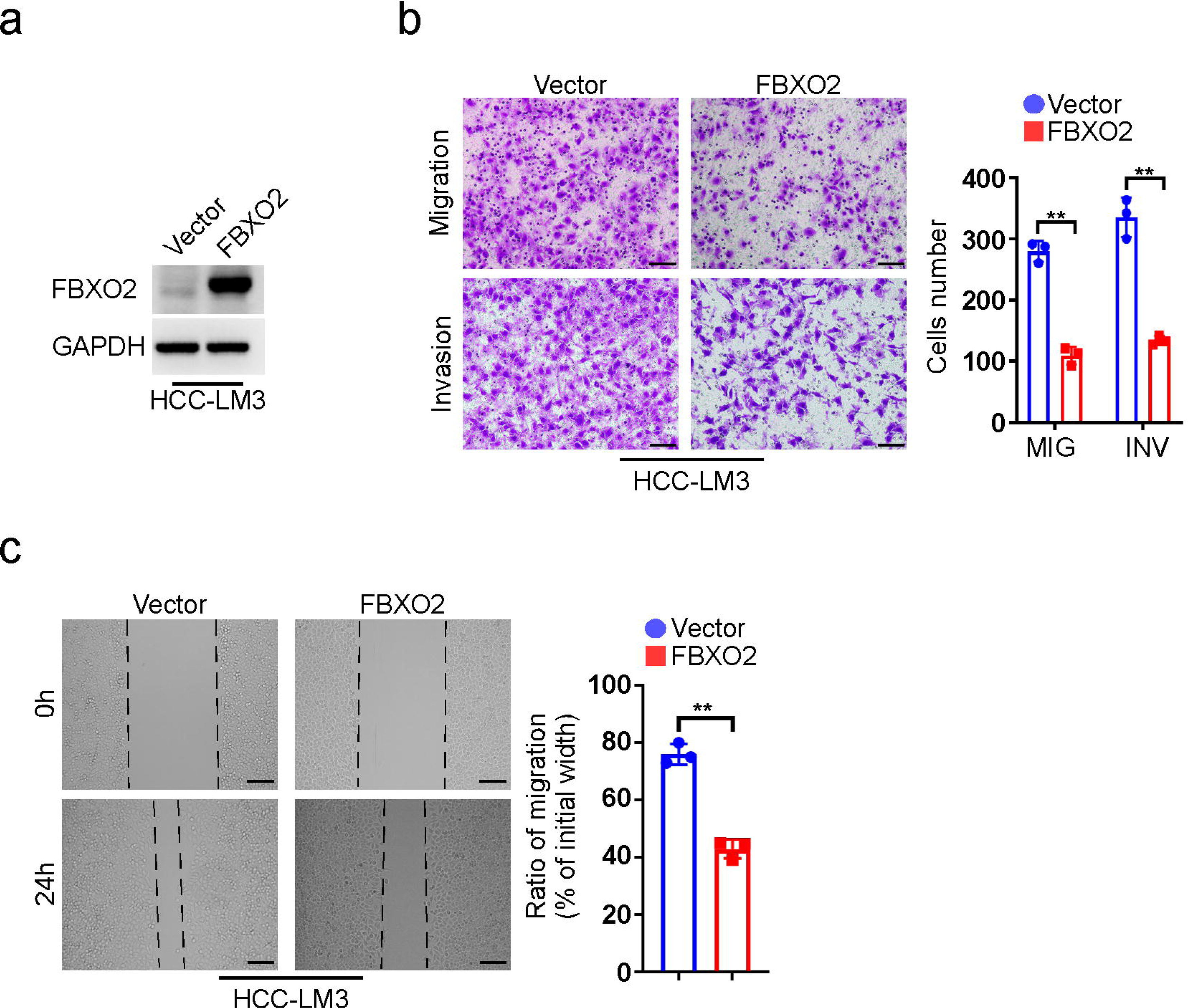
FBXO2 suppresses HCC tumor cells migration and invasion. **a,** IB analysis of FBXO2 protein inHCC-LM3 cells with or without FBXO2 overexpression. **b, c,** FBXO2 overexpression inhibits migration or invasion of HCC-LM3 cells through transwell chamber migration and invasion assays (**b**) and wound healing assay (**c**). Scale bar, 50 μm. Data are the meanL±LSD. Statistical significance is determined by a two-tailed *t* test. \*\**P* < 0.01.

**Extended Data Fig. 3.**
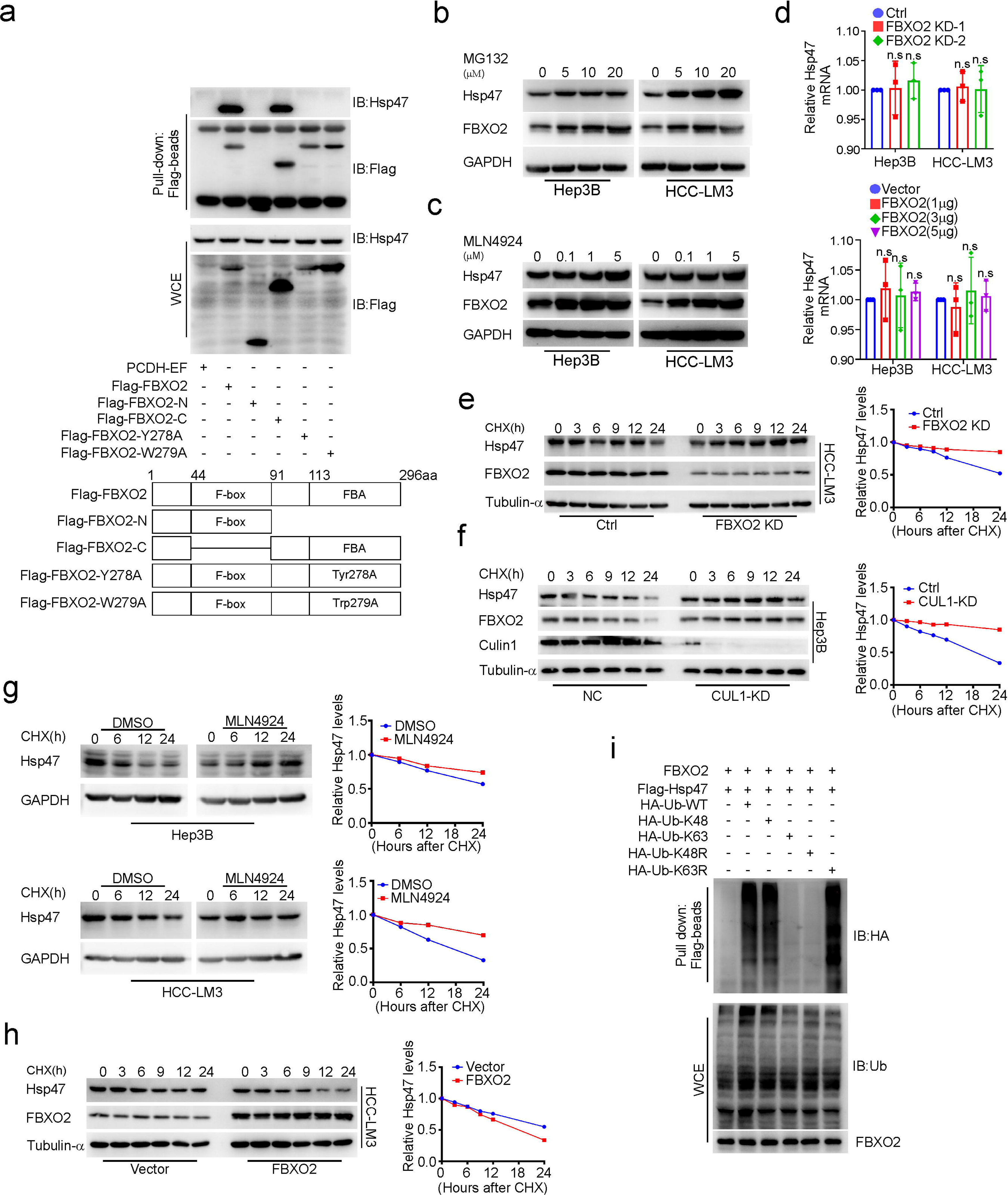
FBXO2 acts as an E3 ligase of Hsp47. **a,** The FBA domain of FBXO2 is responsible for its binding with Hsp47. HEK293 cells were transfected with Flag-tagged WT (Flag-FBXO2) or FBXO2 variants as indicated followed by Flag-beads pull-down. Precipitates and WCE were immunoblotted with the indicated antibodies. **b, c,** IB analysis of the indicated proteins in Hep3B and HCC-LM3 cells treated with different doses of MG132 **(b)** or MLN4924 **(c)** for 12 h. **d**, qRTLPCR analysis of Hsp47 mRNA in FBXO2-KD or -overexpressing Hep3B or HCC-LM3 cells. Data are displayed as the mean ± SD (n=3). **e,** IB analysis of the indicated proteins in HCC-LM3 cells with or without FBXO2 KD followed by cycloheximide (CHX, 50mg ml^-1^) treatment for the indicated time periods. The band intensity was quantified using Image J, normalized to Tubulin-a first, then normalized to the t=0 time point. **f,** CUL1 ablation extends the half-life of Hsp47. IB analysis of the indicated proteins in Hep3B cells with or without CUL1 KD. The treatments of the cells and the quantification of the band intensity were performed as above. **g**, IB analysis of the indicated proteins in Hep3B and HCC-LM3 cells treated with or without MLN4924 (0.1 mM) followed by CHX (50 mg ml^-1^) treatment for the indicated times. The band intensity was quantified using Image J, normalized to GAPDH first, then normalized to the t=0 time point. **h,** IB analysis of the indicated proteins in HCC-LM3 cells with or without overexpression of FBXO2 followed by CHX treatment for the indicated time periods. The band intensity was quantified as **e**. **i,** FBXO2 promotes Hsp47 ubiquitylation via K48 linkage: IB analysis of Flag-beads pull-down and WCE derived from HEK293 cells transfected with the indicated constructs.

**Extended Data Fig. 4.**
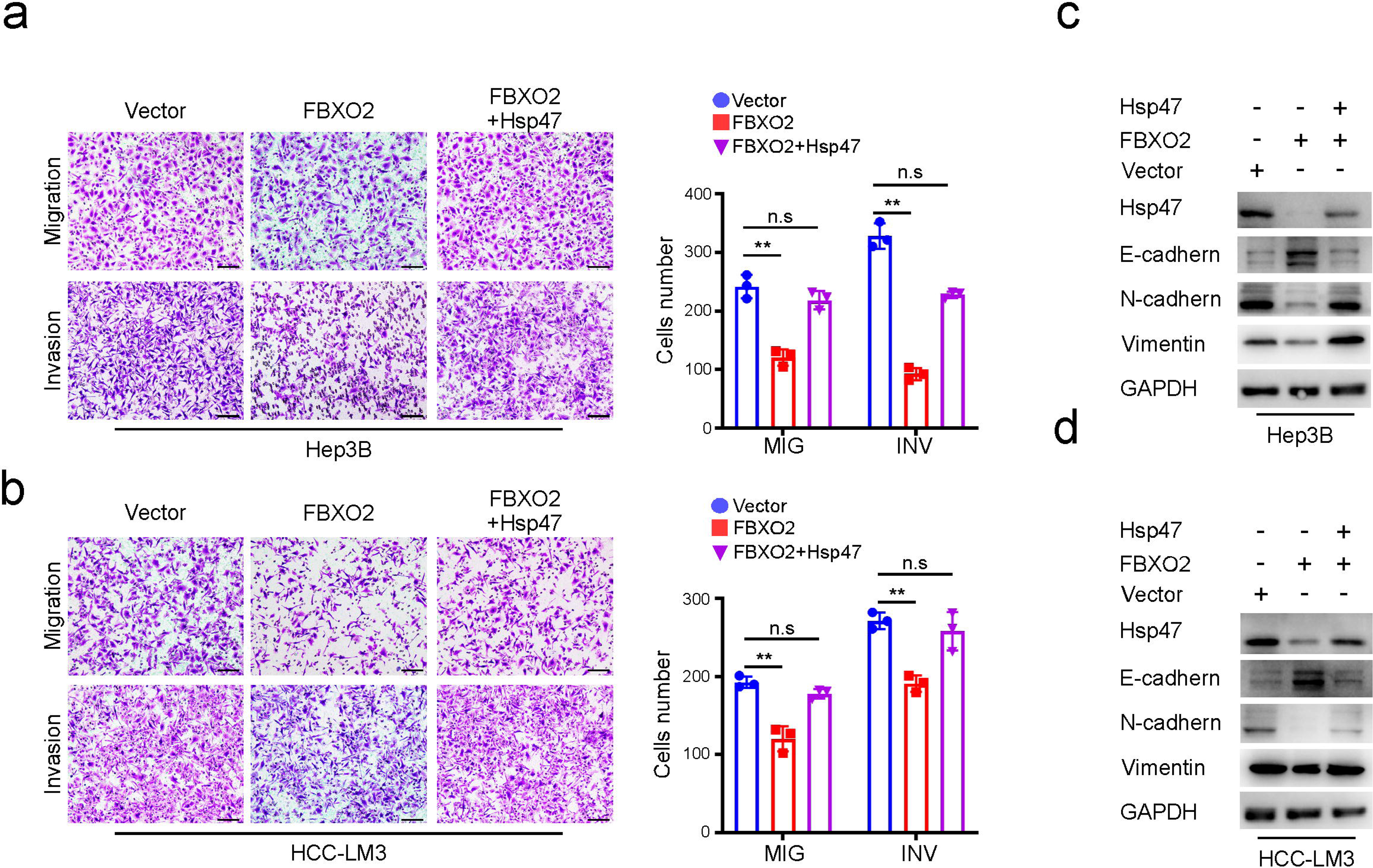
FBXO2 inhibits migration and invasion of HCC tumor cells by down-regulating Hsp47. **a, b,** Representative images and quantification of the effects of overexpression of FBXO2 or FBXO2 plus Hsp47 on the migration and invasion of Hep3B **(a)** and HCC-LM3 **(b)** cells through transwell chamber migration and invasion assays. Scale bar, 50 μm. Data show the meanL±LSD. Statistical significance determined by two-tailed t-test. \*\**P* <0.01. **c, d,** IB analysis of the indicated proteins in above cells.

**Extended Data Fig. 5.**
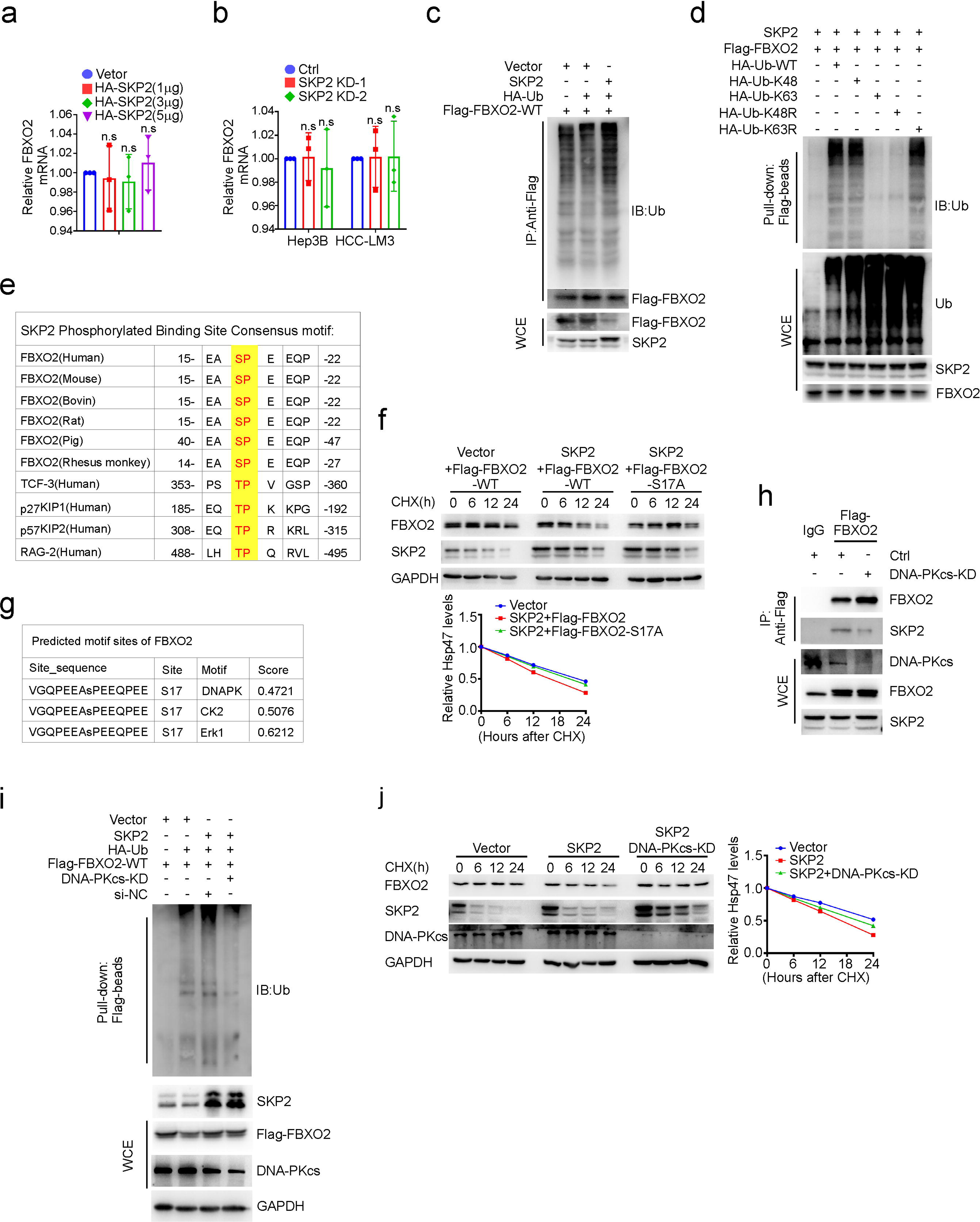
SKP2-mediated polyubiquitination and half-life of FBXO2 depends on DNA-PKcs. **a**, **b**, qRT-PCR analysis of FBXO2 mRNA in SKP2-overexpressing (**a**) or -ablated (**b**) Hep3B and HCC-LM3 cells. Data are shown as the mean ± SD. **c**, SKP2 promotes polyubiquitination of FBXO2. IB analysis of anti-Flag IP and WCE derived from HEK293 cells transfected with the indicated constructs. **d**, SKP2 ubiquitinates FBXO2 via K48 linkage. IB analysis of Flag-beads pull-down and WCE derived from HEK293 cells transfected with the indicated constructs. **e,** Evolutionary conservation of SKP2 binding site in FBXO2 protein sequence of different species. **f**, SKP2 shortens the half-life of FBXO2-WT but not the mutant FBXO2-S17A. IB analysis of the indicated proteins in parental (Vector) or SKP2-overexpressing Hep3B cells transfected with the indicated constructs followed by CHX treatment for the indicated periods. The band intensity was quantified using Image J, normalized to GAPDH, and then normalized to the t=0 time point. **g,** The top three phosphorylation kinases of FBXO2 predicted by SCANSITE 4.0 (https://scansite4.mit.edu/4.0/#home). **h**, DNA-PKcs ablation reduces SKP2-FBXO2 binding. IB analysis of Anti-Flag IP and WCE derived from HEK293 cells with or without DNA-PKcs ablation. **i**, DNA-PKcs ablation weakens SKP2-mediated polyubiquitination of FBXO2. IB analysis of Flag-beads pull-down and WCE derived from the indicated construct-transfected HEK293 cells with or without DNA-PKcs deletion. **j**, DNA-PKcs ablation extends the half-life of FBXO2. IB analysis of the indicated proteins in parental (Vector) or SKP2-overexpressing Hep3B cells with or without DNA-PKcs ablation followed by CHX treatment for the indicated periods. The band intensity was quantified using Image J, normalized to GAPDH, and then normalized to the t=0 time point.

## Supplementary

**Supplementary Table 1 The sequences of siRNAs and shRNAs.**

**Supplementary Table 2 The primer sequences for qRT-PCR.**

**Supplementary Table 3 Antibodies used in this study.**

**Supplementary Table 4 Mass spectrometry results for Co-IP.**

**Supplementary Table 5 The primer sequences for genotyping of mouse.**

