## Supplementary for "SKP2-mediated FBXO2 proteasomal degradation drives hepatocellular carcinoma progression via stabilizing Hsp47"

**Supplementary Table 1. The sequences of siRNAs and shRNAs**

| **shRNA** | **sense（5'-3'）** |
| --- | --- |
| shFBXO2#1: | CCGGGAUGAGAGCGUCAAGAAGUCTCGAGACTTCTTGACGCTCTCATCTTTTTG |
| shFBXO2#2: | CCGGCAGUUCUACUUCCUGAGCACTCGAGTGCTCATTAAGTAGTTCTGTTTTTG |
| shFBXO2#3： | CCGGAAGGUAGAUAGGCCUUAACCTCGAGGTTAAGGCCTATCTACCTTTTTTTG |
| shSKP2#1 | CCGGGATAGTGTCATGCTAAAGAATCTCGAGATTCTTTAGCATGACACTATCTTTTTG |
| shSKP2#2 | CCGGGCCTAAGCTAAATCGAGAGAACTCGAGTTCTCTCGATTTAGCTTAGGCTTTTTG |
| shHsp47#1 | CCGGAGCCCTCTTCTGACACTAACTCGAGTTAGTGTCAGAAGAGGGCTTTTTTG |
| shHsp47#2 | CCGGCCTCTACAACTACTACGACGACTCGAGTCGTCGTAGTAGTTGTAGAGGTTTTTG |
| **siRNA** | **sense（5'-3'）** |
| si-FBXO2#1: | GAUGAGAGCGUCAAGAAGU |
| si-FBXO2#2: | CAGUUCUACUUCCUGAGCA |
| si-FBXO2#3： | AAGGUAGAUAGGCCUUAAC |
| si-SKP2#1 | GATAGTGTCATGCTAAAGAAT |
| si-SKP2#2 | GCCTAAGCTAAATCGAGAGAA |
| si-Hsp47#1 | AGCCCTCTTCTGACACTAA |
| si-Hsp47#2 | CCTCTACAACTACTACGACGA |
| si-DNA-PKcs | GATCGCACCTTACTCTGTT |

**Supplementary Table 2. The primer sequences for qRT-PCR**

| Primer names | Sequences (5'-3') |
| --- | --- |
| H-GAPDH-F | CGCTCTCTGCTCCTCCTGTT |
| H-GAPDH-R | CCATGGTGTCTGAGCGATGT |
| H-FBXO2-F | ACTTGGAAGGCTGGTGTGAC |
| H-FBXO2-R | TCAAAGGAGGAGGCGAAGTA |
| H-Hsp47-F | TGAAGATCTGGATGGGGAAG |
| H-Hsp47-R | CCGCACTAGGAAGATGAAGG |

Abbreviations: H, Human; F: Forward; R: Reverse.

**Supplementary Table 3. Antibodies used in this study**

| **Antibodies** | **Companies** | **Catalog#** | **Application** |
| --- | --- | --- | --- |
| FBXO2 | Proteintech | #14590-1-AP | 1:1000 (WB) 1:200 (IHC) |
| FBXO2 | Santa Cruz | #sc-398111 | 1:1000(WB) |
| Hsp47 | Proteintech | #10875-1-AP | 1:1000 (WB) 1:200 (IHC) |
| SKP2 | Abcam | #ab183039 | 1:1000 (WB) 1:200 (IHC) |
| E-cadhrein | CST | #3195S | 1:1000 (WB) |
| N-cadherin | CST | #13116S | 1:1000 (WB) |
| Vimentin | CST | #5741S | 1:1000 (WB) |
| b-catenin | CST | #8480S | 1:1000 (WB) |
| c-Met | Proteintech | #25869-1-AP | 1:1000 (WB) |
| EGFR | CST | #54359S | 1:1000 (WB) |
| DNA-PKcs | Proteintech | #19983-1-AP | 1:1000 (WB) |
| CK2a | Proteintech | #24883-1-AP | 1:1000 (WB) |
| Ubiquitin | CST | #3936S | 1:1000 (WB) |
| FLAG | Sigma-Aldrich | #F1804 | 1:1000 (WB) |
| HA | Invitrogen | #26183 | 1:1000 (WB) |
| HIS | CST | #12698S | 1:1000 (WB) |
| β-Tubulin | CST | #2146s | 1:1000 (WB) |
| GAPDH | CST | #5174S | 1:1000 (WB) |
| Ki67 | Abcam | #ab15580 | 1:400 (IHC) |
| HRP, Anti- rabbit IgG | CST | #7074 | 1:3000 (WB) |
| HRP, Anti-mouse IgG | CST | #7076 | 1:3000 (WB) |

**Supplementary Table 4. Mass spectrometry results for Co-IP**

| Proteins | ΣCoverage | Σ# Proteins | Σ# Unique Peptides | Σ# Peptides |
| --- | --- | --- | --- | --- |
| FBXO2 | 75.37 | 3 | 15 | 15 |
| TMEM256-PLSCR3 | 68.29 | 3 | 1 | 1 |
| HYOU1 | 62.66 | 27 | 52 | 52 |
| EI24 | 57.14 | 9 | 1 | 1 |
| SERPINH1 | 55.5 | 16 | 19 | 19 |
| RPL38 | 50 | 4 | 5 | 5 |
| EMC4 | 48.61 | 9 | 3 | 3 |
| PLOD3 | 47.43 | 10 | 26 | 26 |
| NOMO3 | 47.38 | 11 | 2 | 42 |
| NOMO1 | 46.97 | 13 | 2 | 42 |
| CSTB | 45.92 | 3 | 3 | 3 |
| ATP5MF-PTCD1 | 44.44 | 6 | 2 | 2 |
| PSMA6 | 43.1 | 6 | 2 | 2 |
| HLA-A | 42.54 | 2932 | 0 | 6 |
| HLA-A | 41.44 | 1648 | 1 | 6 |
| POGLUT3 | 40.04 | 3 | 16 | 16 |
| HLA-B* | 37.13 | 3811 | 0 | 7 |
| HLA-B | 37 | 6791 | 0 | 7 |
| HLA-B | 37 | 3526 | 0 | 7 |
| COLGALT1 | 36.98 | 7 | 21 | 22 |
| TFRC | 36.32 | 6 | 22 | 22 |

**Supplementary Table 5. The primer sequences for genotyping of mouse**

| Primer names | Sequences (5'-3') |
| --- | --- |
| FBXO2-F1 | CTCAGAGCAGTACCTTGCTCATAAG |
| FBXO2-R1 | GCTAGTAGACACTCACAGGAATACGG |
| FBXO2-F2 | GCATCGCATTGTCTGAGTAGGTG |
| FBXO2-R2 | TGTAACTATGGCTGGCCTAGAGC |
| Alb-cre-F | ATTTGCCTGCATTACCGGTC |
| Alb-cre-R | ATCAACGTTTTCTTTTCGG |
| Trp53-F | GGTTAAACCCAGCTTGACCA |
| Trp53-R | GGAGGCAGAGACAGTTGGAG |

Abbreviations: F: Forward; R: Reverse.
